# Charged Scanning Mutagenesis as a High Throughput Approach for Epitope Mapping

**DOI:** 10.1101/2025.07.20.665262

**Authors:** Kawkab Kanjo, Munmun Bhasin, Chandrani Dey, Navid Bakshi, Aparna Ashok, Randhir Singh, Sahil Kumar, Rajesh P. Ringe, Raghavan Varadarajan

**Affiliations:** Molecular Biophysics Unit (MBU), Indian Institute of Science, Bengaluru, 560012, India; Department of Biotechnology, University of Calcutta, Kolkata, West Bengal 700073, India; Department of Biotechnology, University of Kashmir, Srinagar, 190006, India; Mynvax Private Limited; 3rd Floor, Brigade MLR Centre, No.50, Vani Vilas Road, Basavanagudi, Bengaluru, 560004, India; Virology Unit, Institute of Microbial Technology, Council of Scientific and Industrial Research (CSIR), Chandigarh, 160036, India

**Keywords:** Mutagenesis, epitope mapping, library, charged scanning, receptor binding domain

## Abstract

Identifying neutralizing epitopes is important for developing vaccines and inhibitors against viral pathogens. We describe a rapid method for epitope mapping, employing barcoded charged scanning mutagenesis libraries displayed on the yeast surface, and screening using flow cytometry coupled with deep sequencing. Prior scanning mutagenesis data suggest that mutations to a charged residue, such as Aspartic acid or Arginine, will be well tolerated at exposed positions of an antigen, and minimally affect protein stability and expression. Yet such substitutions at epitope residues strongly perturb binding to a cognate partner. We constructed an Aspartate scanning library of SARS-CoV-2 RBD and linked every mutation in the library to a defined unique barcode. The approach was used to map epitopes targeted in polyclonal sera of mice immunized with different SARS-CoV-2 immunogens. In contrast to complete mutational scans, charged scanning mutagenesis with the introduced barcoding strategy employs libraries with >50-fold lower diversity, facilitating library construction, screening, and downstream analysis, and also allowing for further multiplexing of samples, thus accelerating interaction site identification, as well as vaccine and inhibitor development.

## Introduction

Mapping protein-protein interactions (PPIs) and antibody epitopes is a crucial requirement for studying protein function and developing effective interventions for human diseases. Epitopes are a part of an antigen that is recognized by an antibody and can be classified into continuous (linear) or discontinuous (conformational) epitopes ^1^. Mapping linear epitopes is a straightforward process, in contrast to that for conformational epitopes. Yet 90% of the epitopes recognized by antibodies raised against an antigen are conformational ^2^. Knowledge of neutralizing epitopes is a critical demand in vaccine development, particularly in epitope-based vaccine design approaches, where a vaccine is designed by stabilizing and maximizing the exposure of neutralizing epitopes to the immune system. Several methods have been employed for epitope mapping of antibody epitopes and protein-protein interactions (PPIs). X-ray crystallography ^3^ is considered the gold standard, though Cryo-EM is now often employed for this purpose ^4–6^. Nuclear Magnetic Resonance (NMR) ^7,8^ and Hydrogen-Deuterium exchange coupled to Mass Spectrometry (HDX-MS) ^9,10^ are two other powerful structural epitope mapping techniques. However, these methods are time-consuming, labor-intensive, low-throughput and require optimization and highly purified soluble protein samples. Recently, Deep Mutational Scanning (DMS) has been widely used as an alternative functional PPI and epitope mapping approach. In DMS, every residue in the target protein is mutated to all other amino acids, which allows the identification of the interacting or epitope residues as the mutations that impact or escape binding to an interacting partner or an antibody ^11,12^. Construction and characterization of DMS libraries can be laborious and time-consuming, and data analysis is complicated due to the large size and diversity of the DMS libraries. There is a need to develop rapid, simple, and efficient techniques that employ libraries with lower diversity and less complexity than a complete DMS library as a tool for mapping PPI and antibody epitopes.

Cysteine scanning mutagenesis coupled to chemical labelling is an attractive approach for mapping epitopes and binding sites since Cysteines are less frequent in the proteome, well tolerated when introduced at exposed positions, introduce minimum to no structural perturbation and facilitate site-specific protein modification ^13–15^. A Cysteine scanning mutagenesis library can be generated by substituting every residue on the protein surface with cysteine, followed by protein expression and labelling of the surface-exposed cysteines with a cysteine-specific bulky group. This leads to masking the antigen surface and blocking the binding of an interacting partner to its binding site. We previously described an efficient cysteine epitope mapping strategy using yeast surface display coupled with chemical labelling, FACS, and deep sequencing ^15^ and subsequently, extended this to map neutralizing antibodies (NAbs) epitopes against the HIV-1 virus ^16^.

Charged amino acid scanning mutagenesis represents a simpler epitope mapping methodology with potential advantages over other scanning mutagenesis methods ^17^. Charged amino acid scanning can be used to probe structural features of proteins. Charged substitutions are well tolerated at exposed sites with minimum or no effect on protein conformation and expression, while they are poorly tolerated at buried sites, leading to structural perturbation, protein misfolding, and lowered expression, with aspartate having the most destabilizing effect ^17^. Hence, Aspartate can be used as a probe for the identification of buried and exposed residues in proteins, minimizing the need for protein three-dimensional structural information. Substitutions of amino acids at exposed sites in a protein with charged amino acids have a minimal or no impact on protein expression or conformation. However, in case the substituted residues are part of an epitope or interface that interacts with a partner, the binding of the protein with its interacting partner is severely affected. High-throughput and rapid functional probing of protein-protein interactions using charged scanning libraries can be achieved using the yeast surface display platform (YSD) for assessing the binding and expression of all variants displayed on the surface of yeast cells ^18,19^. The interacting residues can be identified using flow cytometry by the loss of binding to their cognate partner with minimal to no perturbation to the protein conformation or expression.

We have developed a charged scanning mutagenesis library of the SARS-CoV-2 RBD. The exposed positions were mutated to aspartate, and wherever there is an aspartate, asparagine, or glutamate, these were mutated to the positively charged amino acid, arginine. The library was barcoded by fusing a 6-nucleotide (6N) barcode of a known sequence and displayed on the yeast surface. This approach, coupled with flow cytometry and deep sequencing, is useful for high-throughput screening and epitope mapping of monoclonal and polyclonal sera.

## Results

### Library construction and yeast surface display

We hypothesized that substitutions of the protein-exposed amino acids with charged amino acids would have minimum or no impact on protein expression and conformation, but would significantly affect the binding of the protein with its interacting partner only if the substituted residues were part of the epitope or interface that interacts with that partner. Since charged amino acids are not tolerated at buried sites ^17^, we expected both the expression and the binding of charged substitutions at these sites to be diminished. To validate the approach described above, we analysed a few previously published datasets of site saturation mutagenesis libraries (Figure 1). Three of the data sets represent deep mutational scanning data that were analysed previously ^20^. These datasets were re-analysed, and each substitution was assigned a score between 0 and −1.0 based on the effect on binding to a cognate partner, where a 0 score means the substitution’s effect on binding is similar to wild type, and a −1.0 score means high sensitivity of the active site to this substitution and severe impact on binding (Figure 1A-C). It is seen from the analysed datasets that charged substitutions have clear discriminating potential between active site and non-active site residues (Figure 1). Charged substitutions at active-site residues severely affect the binding of the proteins to their cognate partners to a resolution sufficient to map these active sites accurately. Starr et al. described a Site Saturation Mutagenesis (SSM) library of the SARS-CoV-2 receptor binding domain (RBD) displayed on the yeast surface, where the expression and binding of all variants to Angiotensin Converting Enzyme-2 (ACE2) receptor were characterized ^21^. A subset of the SSM library that contains only charged amino acid substitutions at the ACE2 binding site was selected and analysed. The ACE2 binding was severely impacted upon the substitution of binding site residues with charged amino acids (D, E, K, R) (Figure 1D), and the highest impact was observed for substitution to Aspartate. Figure 1E shows the Box plot distribution of mutational sensitivity scores for charged substitutions. The medians (Middle Lines of the Boxes) for D and E are positioned lower on the ‘Mutational sensitivity score’ axis compared to K and R. This suggests that the typical (median) mutational sensitivity scores for groups D and E are lower (more negative) than for groups K and R. Specifically, Group D’s median (-0.65) is the lowest among all groups, followed closely by Group E (-0.6).

**Figure 1.**
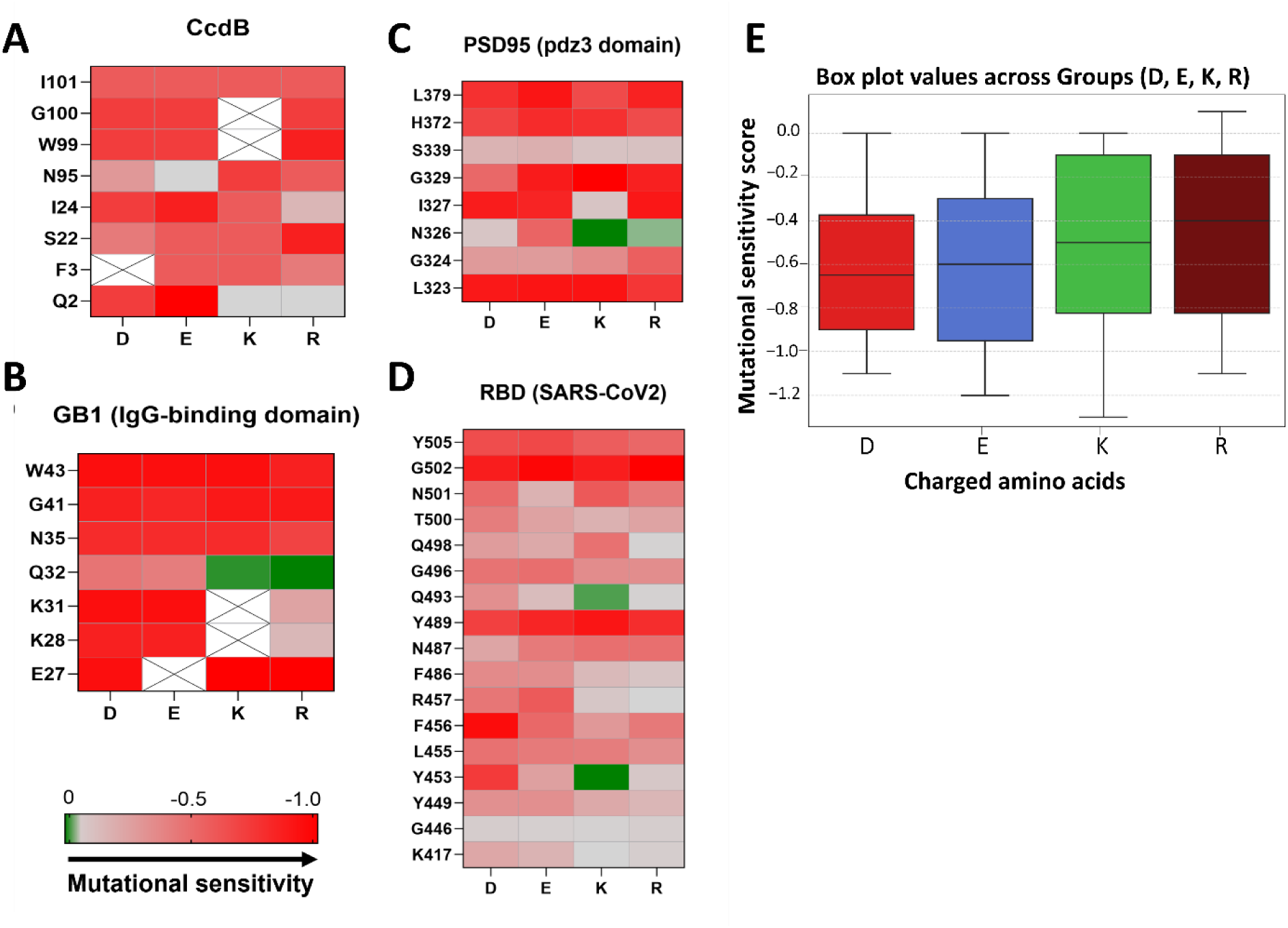
Mutational effects inferred from deep sequencing of charged substitutions at active-site residues involved in protein:protein interactions. The horizontal and vertical axes represent the active-site residues and charged substitutions, respectively. The color of each heatmap corresponds to the rescaled mutational effect score of the mutant, derived from published data ^20^. The colour gradient ranges from green to red, indicating decreasing mutational effect scores and increasing mutational sensitivity. Mutants with scores close to zero behave similarly to the wild type. Substitutions that were not sequenced in the input or selected pools or that were eliminated during quality filtration steps are depicted in an empty box with a cross. Active site residues were assigned based on crystal or cryo-EM structures of the protein:protein complex. The heatmaps show the charged substitutions within the 4 large-scale mutagenesis datasets utilized. (A) CcdB. (B) GB1 (IgG-binding domain). (C) PSD (pdz3 domain). (D) RBD (SARS-CoV-2). (E) Box plot showing the distribution of mutational sensitivity scores (y-axis) for charged substitutions DEKR (x-axis). Groups are arranged by their observed median score, from most negative to least negative: D, followed by E, K, and R. Within each box, the central line indicates the median. The box represents the interquartile range (IQR), spanning from the first quartile (Q1) at the bottom to the third quartile (Q3) at the top, and includes the middle 50% of the data.

The mammalian codon-optimized SARS-CoV-2 RBD gene was used as a candidate for testing the proposed epitope mapping approach. The Yeast Surface Display (YSD) platform (Figure 2A), coupled with flow cytometry, was used for high-throughput screening. The flow cytometry expression and binding profiles for charged substitutions at the exposed non-epitope residues are expected to be similar to Wild Type (WT). In contrast, such substitutions at exposed epitope residues are expected to maintain similar expression profiles to WT but result in reduced binding (Figure 2B). Introducing charged mutations at buried positions is expected to reduce both the expression and binding of these mutants compared to WT (Figure 2B). Before constructing the library, a few solvent-exposed ACE2 interacting and non-interacting residues and a few buried residues were selected for validation, and charged substitutions were introduced at these positions (Figure 2C). Flow cytometry analysis confirmed the above expectations (Figure 2D-E).

**Figure 2.**
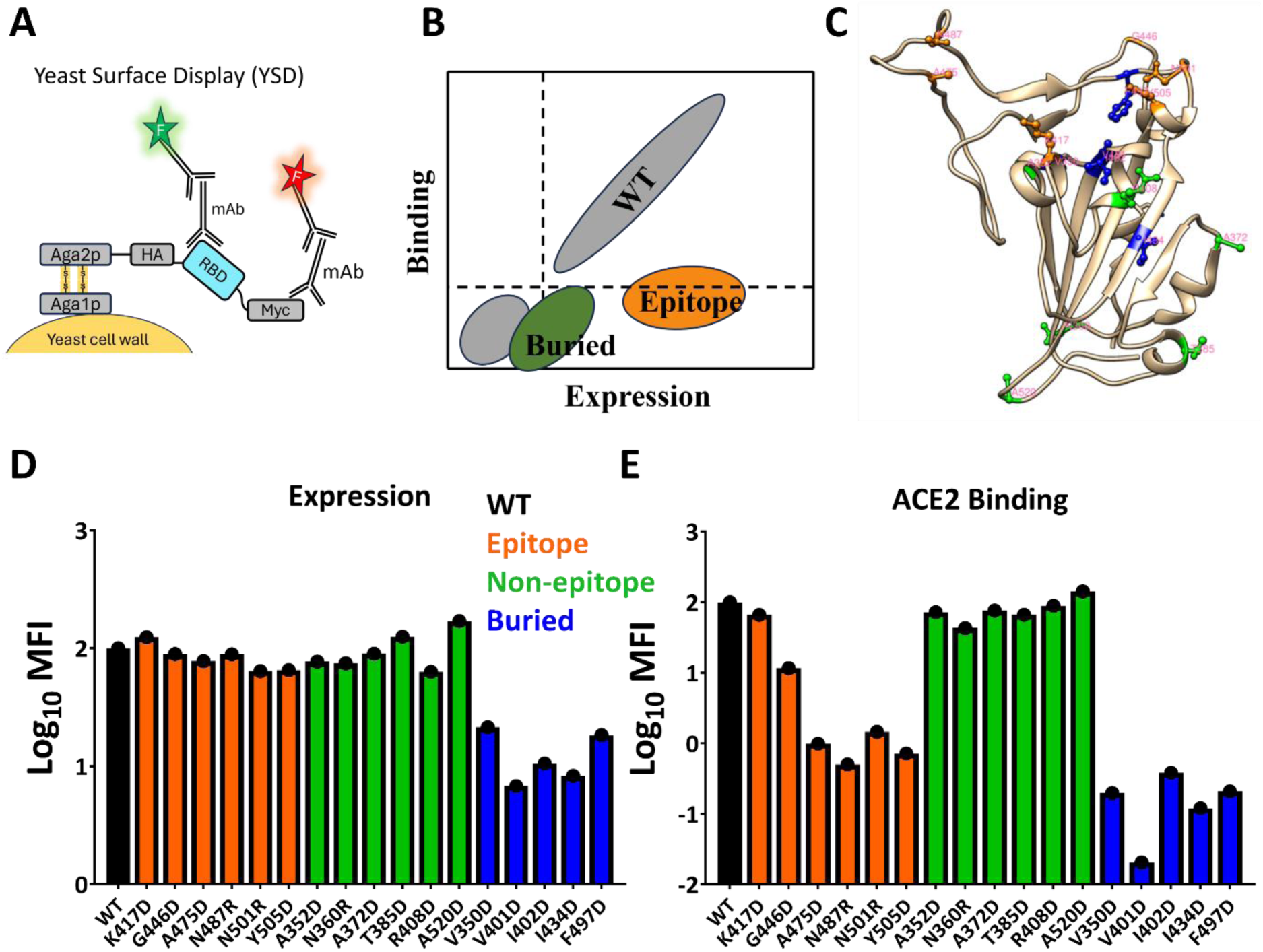
Validation of the epitope mapping approach using charged scanning mutagenesis. (A) A schematic representation of the Yeast Surface Display system. (B) Schematic representation of the expected effect on the expression and binding profiles of the charged amino acid substitutions. (C) Structure of the SARS-CoV-2 RBD with the residues selected for validation displayed in ball and stick, and coloured according to their accessibility and function, as also represented in (D) and (E) (PDB: 6M0J). The yeast surface display expression (D) and ACE2 binding (E) profiles of wild type (black), six solvent exposed and part of ACE2 epitope residues (orange), six solvent exposed and outside ACE2 epitope residues (green), and five buried residues (blue).

The charged scanning library of all the exposed positions of RBD was then constructed, as explained in Figure 3A-B. With a library diversity of 124 barcoded variants (Table S1), tiled sequencing was carried out as shown in Figure 4A, and the reads from individual fragments were analysed and quality filtered with a Phred score > 20 (Figure 4B). Around 48% of the reads passed the quality threshold. The barcodes extracted from the reverse reads were cross-referenced with the pool of barcodes employed during library preparation to identify matches. These matched barcodes were subsequently mapped to the corresponding mutants. This resulted in a higher distribution of the WT reads than the mutant reads across the six fragments, which is expected due to the library’s low diversity, such that the ratio of the fragments containing mutations to the WT fragment is 1:5 (Figure 4C-D). The read analysis confirmed the correct attachment of the 6N barcodes with their expected variants, and barcodes associated with each of the 124 variants were confirmed (Table S2). Interestingly, no wild type was observed in the library, which supports the efficiency of the proposed library construction methodology.

**Figure 3.**
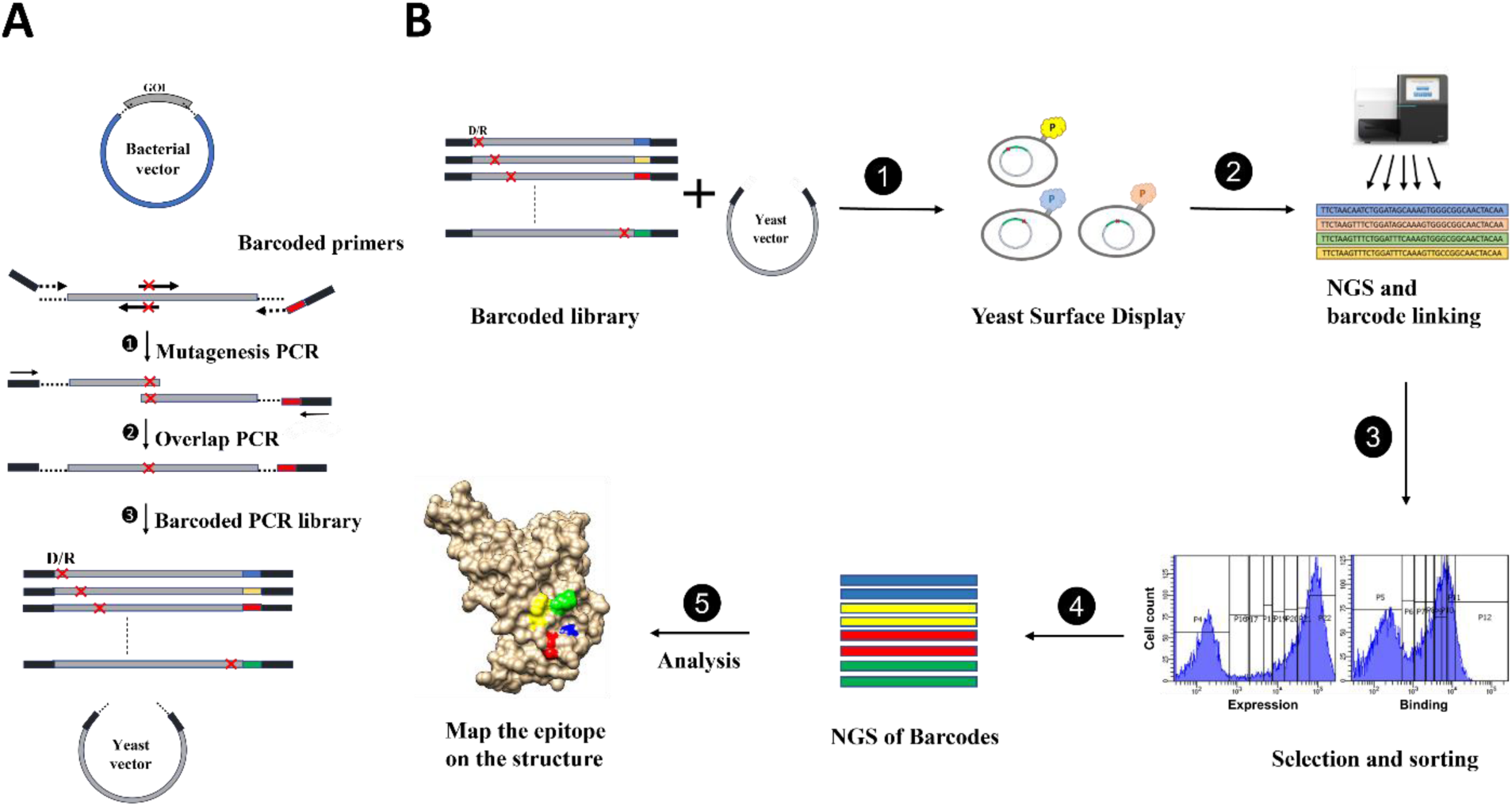
Mutagenesis library construction and barcoding workflow. (A) Mutagenesis by overlap PCR using the WT-RBD cloned in pUC57, a bacterial vector, as a template and employing a common vector-specific forward primer and three position-specific primers: two mutagenesis primers and a vector-specific barcoding reverse primer. The barcodes are introduced to each position in the first round (1) of PCR, and both vector-specific primers are flanked by a yeast-vector overlapping region that is absent in the bacterial vector. The second round (2) of PCR employs two common primers binding to the flanking region of the previous vector-specific primers, hence preventing WT amplification. (B) The workflow for epitope mapping starts from (1) pooling the barcoded joined PCR products from the second round of PCR from the previous step in equimolar ratio and transforming the pooled library along with the double-digested yeast vector into yeast. The transformed cells are plated to estimate the library diversity. The resulting pool can be stored and also subjected to yeast surface display. (2) Deep sequencing of the displayed library to confirm the correct attachment of the desired barcodes with their target mutations. (3) Library subjected to binding with monoclonal antibodies or polyclonal sera and sorting of the cell populations into different bins based on the binding or expression MFI. (4) Deep sequencing of the barcode region from the sorted populations. (5) Analysis of the deep sequencing results and mapping of the identified epitopes on the structure.

**Figure 4.**
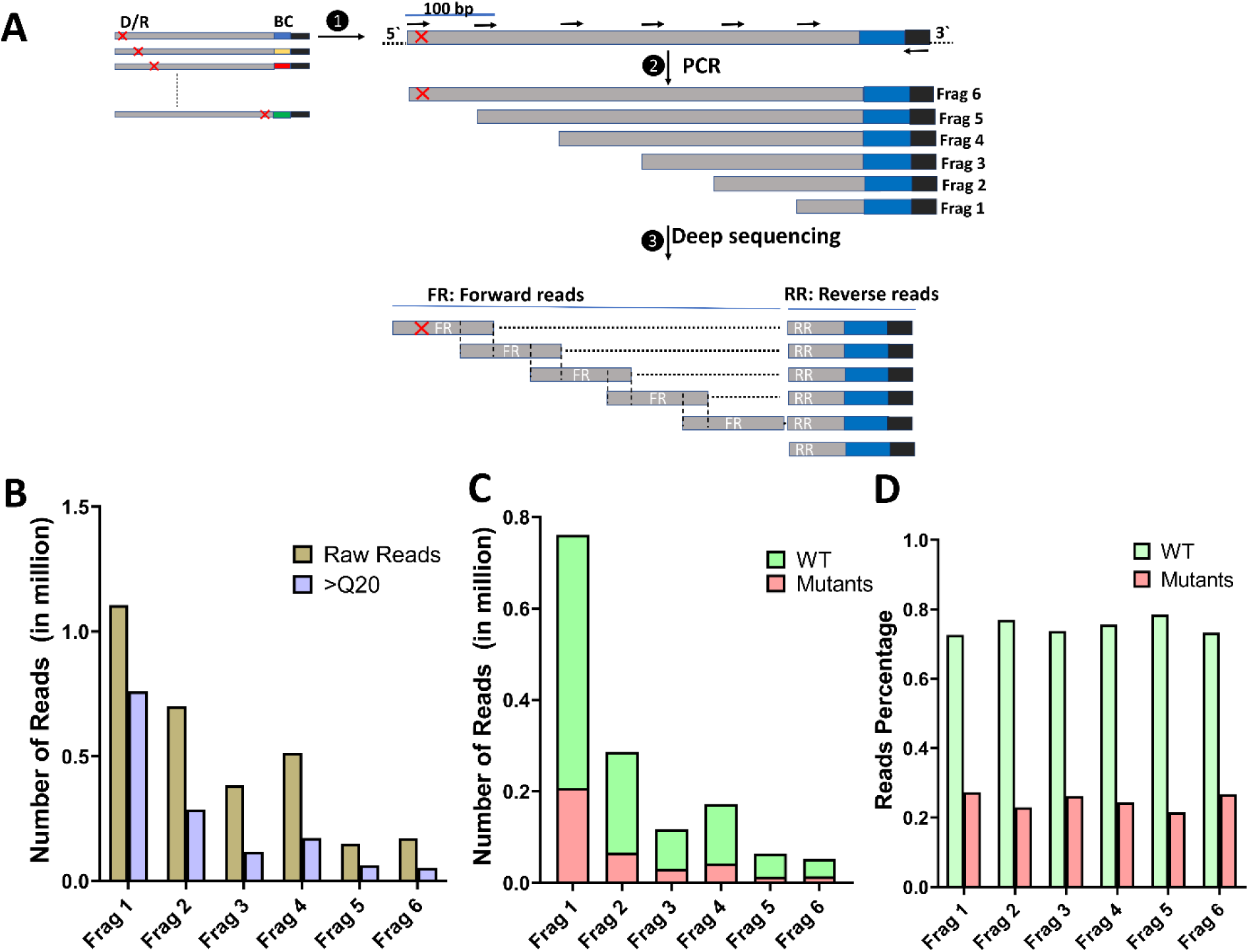
Deep sequencing strategy and linking barcodes to their respective mutations. (A) The plasmids isolated from the YSD library were used as a template for PCR (1). A common reverse primer (black) binds downstream of the barcode region (blue), and a set of forward primers spanning the whole sequence of the gene of interest, with a 100 bp separation between each consecutive primer, was designed. Each forward primer was used in combination with the common reverse primer in separate PCRs to yield different-length PCR products ranging in length from the full length of the gene to 250 bp (2). These PCR products were pooled in equimolar ratio and submitted for Illumina deep sequencing on the NovaSeq 150 PE platform. Each fragment is read for 150 bp from each end to generate, for a given mutant, forward reads from different positions of the gene of interest and reverse reads with an identical barcode for all the fragments generated from the same mutant(3). (B) Raw and quality filtered read distribution in each fragment of the aspartate-barcoded library. The brown bars represent the raw read counts obtained from deep sequencing, while the purple bars depict the subset of reads that passed the quality filter threshold (Q20 cut-off) in each of the six fragments. The number of reads is represented on the y-axis, while each distinct fragment within the library is represented by a bar on the x-axis. (C) Total reads distribution after quality filtering for the wild-type (wt) and mutant sequences. The bar graph depicts the proportion of high-quality reads for the wild-type and mutants, coloured in green and pink, respectively. (D) Percent distribution of WT and mutant sequences for each fragment. Due to the small size of the library and its low diversity, the ratio of WT to mutant sequence is expected to be 5:1 for each fragment, which is close to the observed value.

### Mapping the interaction interface of ACE2 and calculating the variant MFI ratio

The library covers most of the RBD surface, and mutations at most of the reported epitopes of neutralizing antibodies from different classes were included in the library (Figure 5A). To validate and test the efficiency of charged scanning mutagenesis, we mapped the interaction of the RBD with the ACE2 receptor. WT RBD expressed well on the yeast surface and bound to ACE2. When the library was tested for binding to ACE2, a similar profile to WT RBD was observed, except for enrichment in the bins with reduced binding represented by P5, P6, P7, and P8 (Figure 5B). We expect the RBD interacting residues with ACE2 to fall in these bins. The mutants with MFI_Ratio_ one standard deviation higher than the mean MFI were considered interacting or epitope residues. Different values of the cut-off were tested by varying the standard deviation (SD) from 0.1 SD up to 2.5 SD, and the precision and recall were calculated for each SD value tested. We observed a trade-off between precision and recall. The best precision and recall values of 93% and 65%, respectively, were obtained with the 1 SD cutoff value (Figure S1).

**Figure 5.**
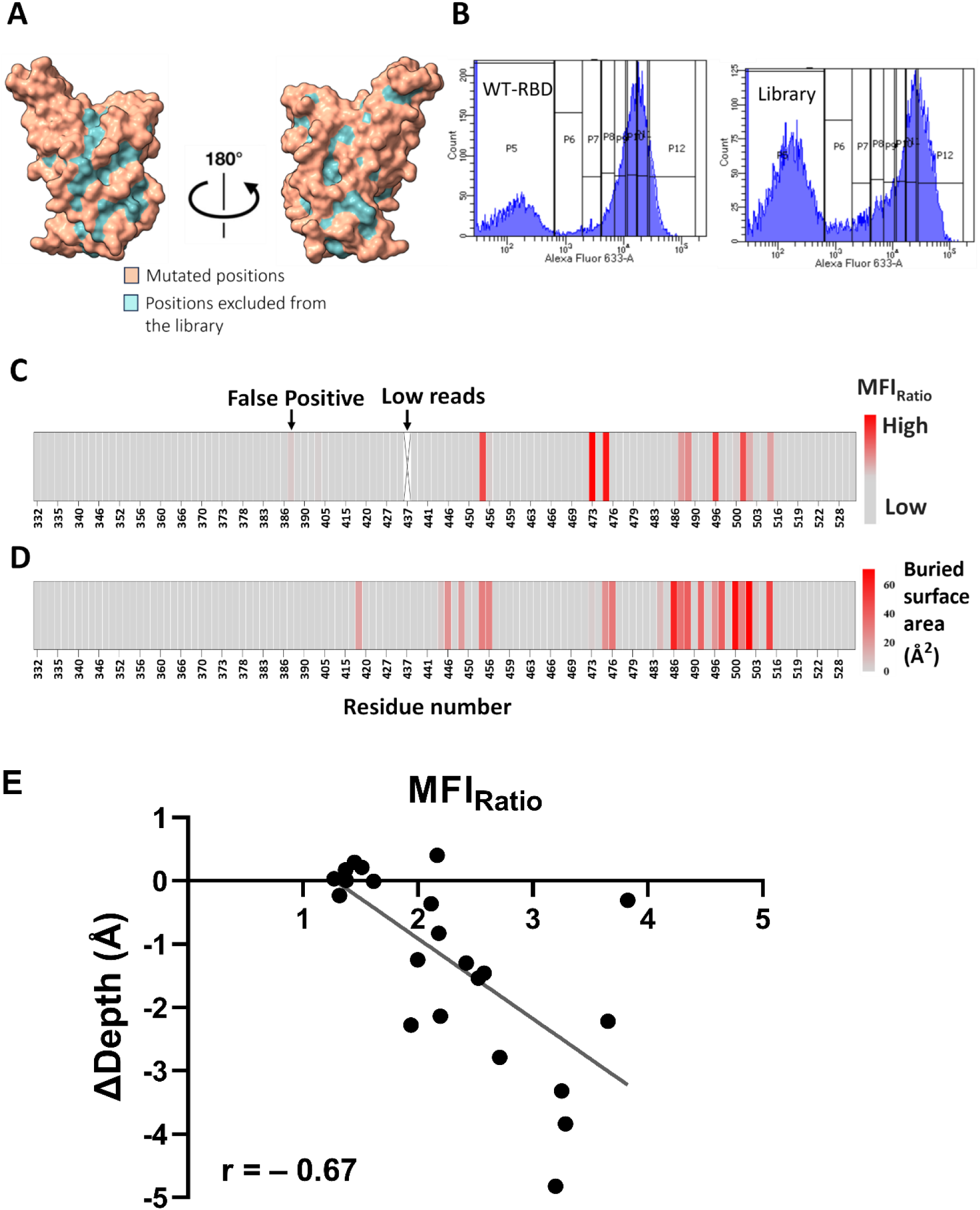
Methodology validation using ACE2 epitope mapping. (A) The mutated positions included in the constructed library with accessibility ≥15% are highlighted on the RBD surface representation in salmon, and the rest of the positions are coloured in cyan (PDB: 6M0J). (B) Flow cytometry histograms of the ACE2 binding profile to WT-RBD and to the charged scanning library. The library population was sorted into eight vertical bins (P5-P12), with bin 5 containing unstained cells and bin 12 containing cells with the maximum binding MFI. (C) The heat map represents the MFI_Ratio_ of the ACE2 sorted bins. Grey colour represents mutants with WT-like MFI_Ratio_ while red represents mutants with higher MFI_Ratio_, the latter are likely epitope residues. From all the epitope residues identified through their MFI_Ratio_ values, we found only one false positive residue. (D) Heatmap showing the change in solvent-accessible surface area upon binding to ACE2. The accessible surface area for residue side chains was calculated from crystal structures of RBD-ACE2 complex (6M0J) using NACCESS V 2.1.1 ^22^. The buried interface surface area was obtained by subtracting the accessible surface area of RBD residue side chains in the presence of ACE2 (PDB ID: 6VXX) from the corresponding area in the absence of ACE2 (PDB ID: 6M0J). (E) Correlation between the MFI_Ratio_ with ΔDepth for all RBD-ACE2 interacting residues, where ΔDepth = Depth (free) – Depth (bound) for each residue.

The MFI_Ratio_ of ACE2 binding for all the library residues was plotted as a heatmap showing the identified interacting residues with a higher MFI_Ratio_ than the rest of the residues in the library in red colour; the darker the red colour, the higher the escape from ACE2 binding exhibited by this mutant (Figure 5C). The identified interacting residues in our analysis were compared with the actual interacting residues, extracted from the ΔASA calculated from the crystal structures (Figure 5D and Table S3). All the residues identified in our scan, except for one, were indeed interacting residues as reported in the literature and confirmed by examining the RBD-ACE2 bound crystal structure. Using this analysis, we were able to recal 65% of the RBD interacting residues with ACE2. When we looked at the MFI_Ratio_ of the remaining interacting residues that were missed in our analysis, these residues had an MFI_Ratio_ that was slightly lower than that of the cutoff MFI_Ratio_, as in the case of the F486D mutation, its MFI_Ratio_ was 0.01 lower than the cutoff. F486D mutation did not show a drastic effect on ACE2 binding, and it has good expression, which explains why its MFI_Ratio_ failed to pass the cutoff for epitope residues. The same applies to residues K417D, V445D, G446D, Y449D, G476D, E484R, Q493D, Q498D, and V503D. The MFI_Ratio_ correlated with change in residue depth upon complex formation (Pearson correlation, r = 0.67, *p* value < 0.001) (Figure 5F). Given the high precision and recall values of 93% and 65%, respectively, as well as the good correlation with residue burial upon complex formation, we next used the proposed epitope mapping methodology to map epitopes from polyclonal sera.

### Epitope mapping of BALB/c mice sera

A set of three sera was obtained from BALB/c mice primed and boosted with 20 µg of either Spike (B.1-S) or RBD subunit vaccines derived from either the ancestral SARS-CoV-2 (B.1-R) or its descendant variant, Omicron (BA.1-R) (Figure 6A). SWE adjuvant was used in all the immunization studies. All the tested sera showed high binding GMT titres of anti-RBD and anti-S6P (six-prolines-stabilized spike) IgG antibodies as determined by ELISA (Figure 6A). The neutralizing activity of the sera was tested using a pseudoviral neutralization assay, and the sera from immunization with B.1 spike or RBD showed high neutralization IC_50_ against B.1 pseudovirus but lower IC_50_ against BA.1 pseudovirus. The sera obtained from immunization with BA.1 RBD neutralized BA.1 pseudovirus with a higher neutralization IC_50_ than the corresponding B.1 IC_50_ (Figure 6A). The known classes of neutralizing epitopes are highlighted on the structure of SARS-CoV-2 RBD and shown in Figure S2. The epitope mapping results from these sera were plotted as a heatmap of the MFI_Ratio_ (Figure 6B). The heatmap suggested that various epitopes in different RBD regions are targeted by these sera. Interestingly, all the sera exhibited similar profiles of the targeted epitopes, whether the immunogen was from an ancestral or Omicron origin.

**Figure 6.**
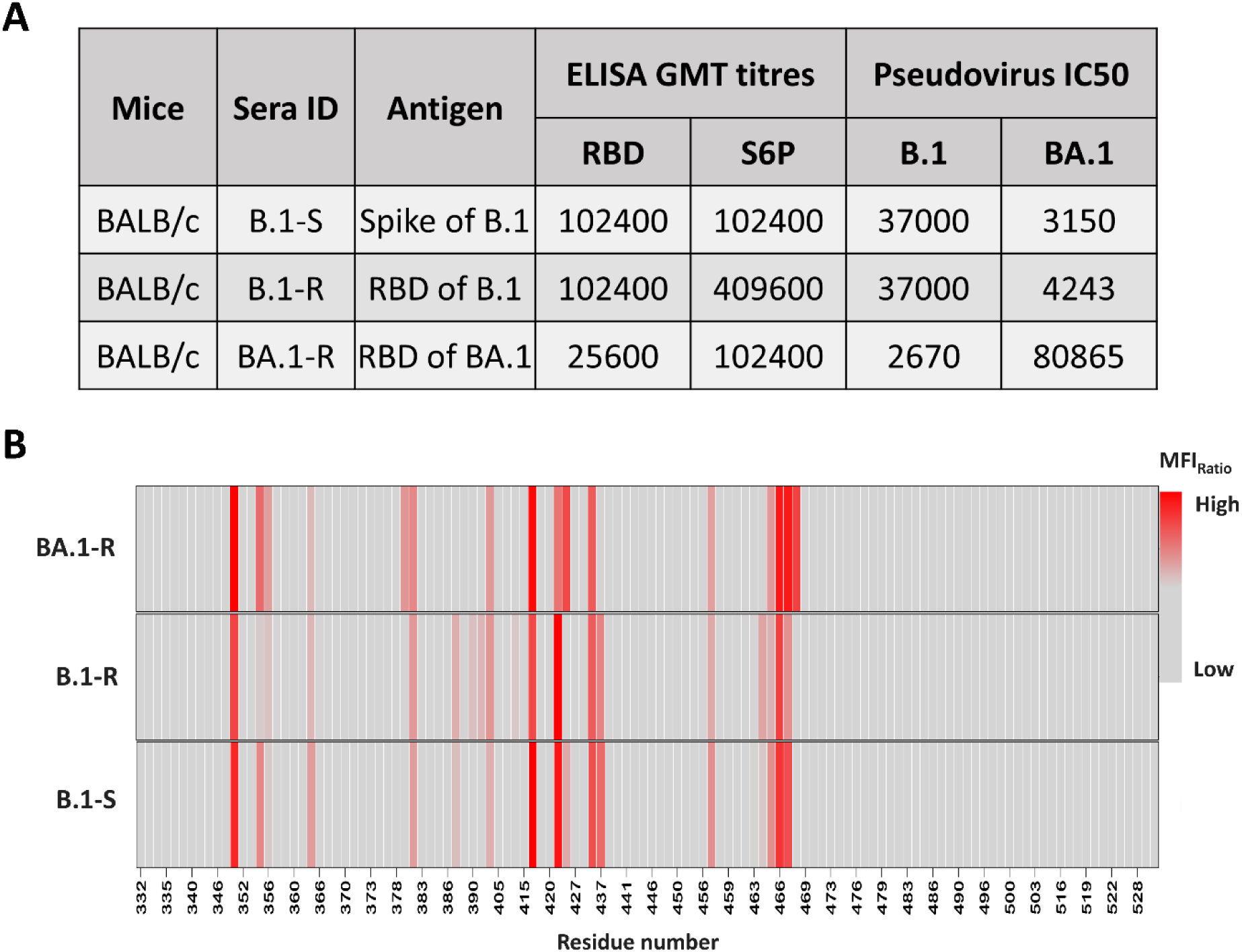
Epitope mapping of polyclonal sera from BALB/c mice. (A) Sera IDs, the corresponding antigen used for immunization, ELISA GMT titres for the individual mice sera tested against RBD and spike S6P protein, and Pseudoviral neutralization IC_50_ values against WT and Omicron BA.1 pseudovirus for the mice sera used in this study. (B) MFI_Ratio_ values. Increasing values of MFI_Ratio_ (red) represent a greater loss of binding upon mutation, indicative of an epitopic residue.

The epitopes were analysed and are highlighted on the RBD structure in Figure 7. Different classes of neutralizing antibody epitopes were targeted by the sera. The most targeted epitope was class I, which overlaps with the ACE2 receptor, followed by class IV and, to a lesser extent, class III. Our analysis also revealed a prominent conserved region targeted by all the tested sera. The region belongs to class V epitopes, a conserved class of neutralizing antibodies that target a rare and cryptic epitope of RBD that lies on the opposite face of the RBM. This epitope was reported in the recent literature, and its epitope residues are highly conserved among all the known VOCs ^5,23^. We also showed in a recent study that the same epitope was targeted in breakthrough infection sera from Indian vaccinees ^24^.

**Figure 7.**
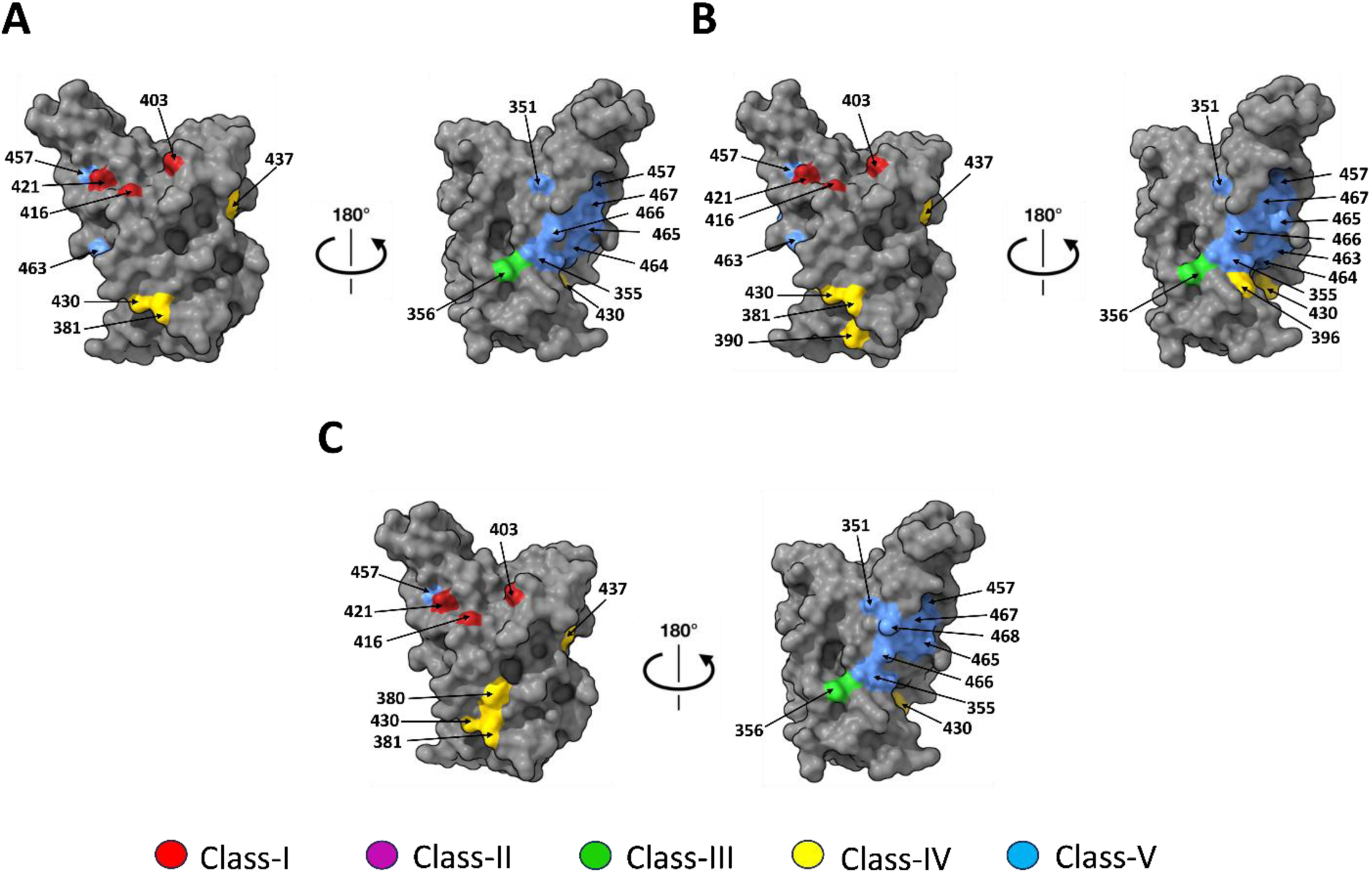
Epitope residues in BALB/c mice polyclonal sera mapped onto the structure of RBD. Putative epitope residues identified from charge scanning mutagenesis are highlighted on the RBD structure (PDB: 6M0J) and coloured according to the class of neutralizing epitope. The class IV and V epitopes are cryptic epitopes that are occluded in the “down” conformation of the RBD in the context of the spike. (A) Serum B.1-S epitopes. (B) Serum B.1-R epitopes. (C) Serum BA.1-R epitopes.

### Epitope mapping of sera from immunized K18-hACE2 transgenic mice

The K18-hACE2 mouse is a transgenic mouse model that expresses the hACE2 receptor on its airway epithelial tissues and hence recapitulates the pathology of human COVID-19 disease ^25^. To compare the humoral immune response elicited in BALB/c to that of K-18-hACE2 mice, we evaluated the antibody response of two sera obtained from K-18-hACE2 mice immunized with either the ancestral (B.1-R) or Omicron (BA.1-R) SARS-CoV-2 RBD (Figure 8A). The anti-RBD and anti-S6P IgG binding GMT titres of B.1-R serum from the transgenic mice were similar to those of the sera from BALB/c mice, but the BA.1-R serum exhibited reduced IgG binding GMT titres against both ancestral RBD and S6P (Figure 8A). The neutralization profile resembles those of BALB/c sera, with the B.1-R sera neutralizing the ancestral pseudovirus better than BA.1, while the BA.1 serum neutralizes BA.1 pseudovirus better than B.1 (Figure 8A). Epitope mapping from K-18-hACE2 transgenic immunized mice sera revealed interesting findings. The identified epitopes targeted by the B.1 serum share some similar residues with epitopes targeted by BALB/c mice sera, and the major epitopes targeted in both belong to class I, class IV, and class V. However, the overall epitope mapping profile of K-18 mice serum differs from that of the BALB/c, such that fewer class V epitope residues were identified in K-18 serum than in BALB/c sera (Figure 8B-D). The serum obtained from K18-hACE2 mice immunized with BA.1 RBD exhibited a completely different profile. The epitopes targeted were variant-specific regions, and most of the escape was focused on the ACE2 binding region (Figure 8B-D). In the Omicron immunized K18-hACE2 mice, the immune response escape was focused on class 1 and class 2, with a smaller amount of class 3 residues. Since the library is made in the WT-RBD background, the epitope residues readily detected are conserved between WT and BA.1 variants.

**Figure 8.**
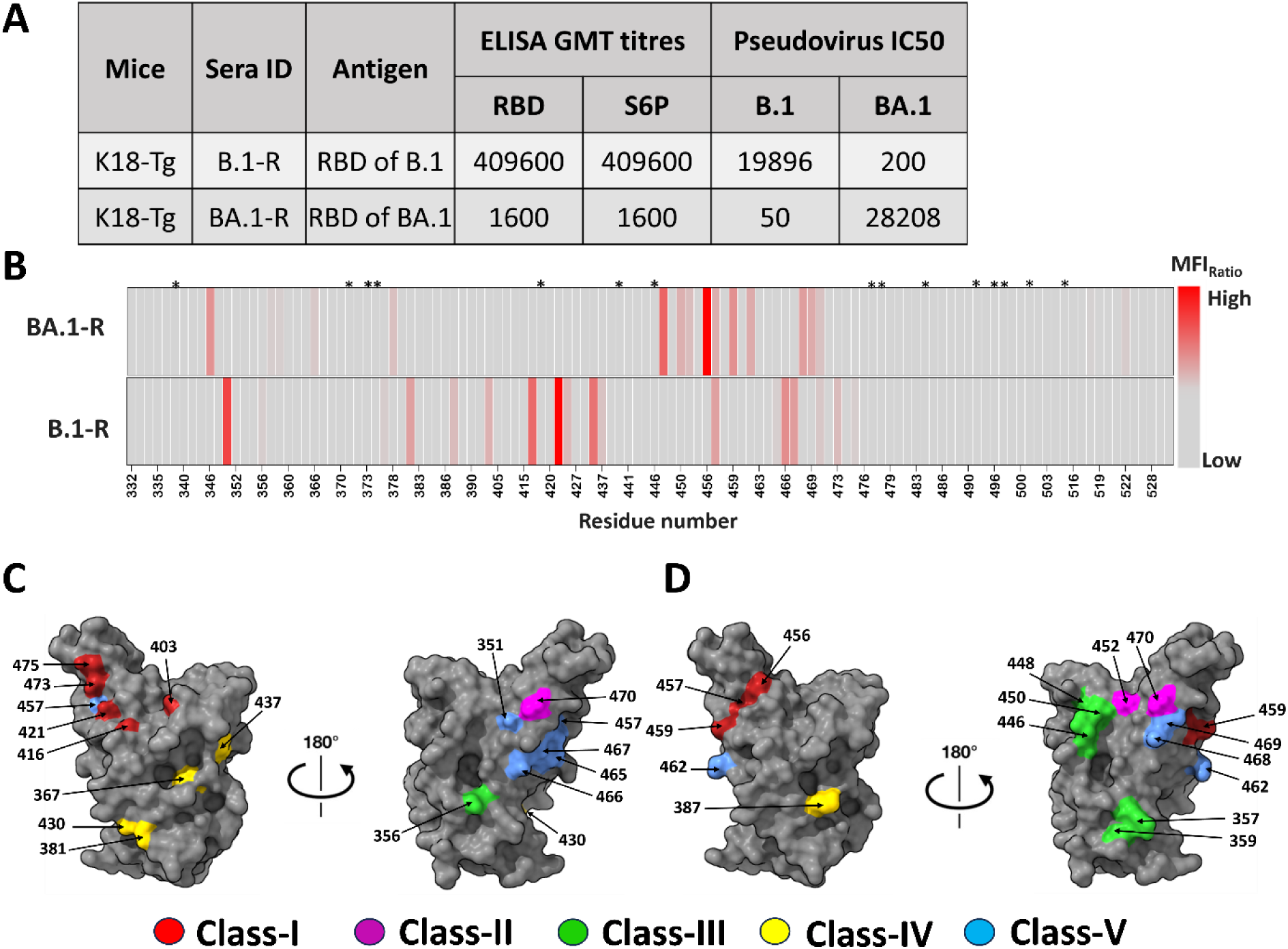
Epitope mapping of polyclonal sera from K18-hACE2 mice. (A) Sera IDs and the corresponding antigen used for immunization. (B) MFI_Ratio_ values. Increasing values of MFI_Ratio_ correspond to an epitope residue (red). Mutated positions in Omicron BA.1 RBD are marked by a *star*. (C-D) Putative epitope residues identified from charge scanning mutagenesis are highlighted on the RBD structure (PDB: 6M0J) and coloured according to the class of neutralizing epitopes. The class IV and V epitopes are cryptic epitopes that are occluded in the “down” conformation of the RBD in the context of the spike. (C) Serum B.1-R epitopes. (D) Serum BA.1-R epitopes.

## 4. Discussion

An infection with a pathogen triggers the humoral and cellular immune responses. Neutralizing antibodies (NAbs) are considered crucial correlates of protection ^26,27^. Epitope-based vaccines aim to elicit high titres of protective neutralizing antibodies ^28,29^. Several epitope mapping techniques were used to probe neutralizing epitopes and identify novel epitopes as a target for vaccine design strategies to achieve this goal.

Deep mutational scanning (DMS) is a powerful tool for studying residue-specific contributions to protein stability and function ^30–32^. In a complete DMS library, every residue is individually randomized. In contrast, the charged scanning mutagenesis approach described here employs much smaller libraries that are easier to construct, screen and analyse because of their limited diversity. Such libraries can be easily utilized along with yeast surface display or other display platforms and flow cytometry for screening. Analysis of published saturation mutagenesis datasets ^20,33^ showed that charged substitutions at active site residues significantly impact protein-protein or protein-antibody interactions.

In this work, we successfully constructed a charged scanning library of all the exposed sites in SARS-CoV-2 RBD. Using the described PCR strategy we proposed, the library included the desired mutated positions with no observations of wild type. The barcoding of the library was done directly through the initial mutagenesis PCR steps using a reverse primer with a known 6-nucleotide sequence unique for each position. This barcoding method is feasible in the case of small-sized and low-diversity libraries. It has an advantage over the random nucleotide barcoding method as it eliminates the requirement for expensive long-read sequencing to link each barcode with its mutation, and Illumina sequencing can be used instead to fulfil the same purpose. The small library size and the suggested barcoding and sequencing strategy render this method suitable for adaptation in low-resource settings, particularly when rapid scanning in a cost-effective manner is a critical demand. The constructed barcoded library was validated by mapping RBD interactions with its receptor, ACE2, with 93% precision and 65% recall of interacting residues. These results are encouraging, especially when achieved by a small-sized library with limited diversity. Our charged scanning results strongly agree with the full DMS scan data, where substitutions of the ACE2 interacting residues to charged residues significantly affected ACE2 binding affinity. Some ACE2 contact positions exhibited plasticity, and mutations were tolerated at these positions, which in turn sheds light on why our approach failed to identify the remaining 35% of the ACE2 interacting residues ^21^. In contrast to earlier published DMS scans ^21,34^, where epitope residues are identified through near complete loss of binding in a 2D histogram, in the present approach, the epitope and interacting residues are identified through sorting of the 1D histograms of the expression and binding cell populations. This provides a semi-quantitative measure of the contribution of each residue to the interaction with the binding partner as can be seen from the data in Figure 5E.

Epitope mapping for monoclonal antibodies is relatively straightforward compared to mapping polyclonal sera containing multiple antibodies targeting different epitopes. Polyclonal sera from BALB/c mice immunized with different SARS-CoV-2 subunit vaccine candidates were subjected to epitope mapping using the charged scanning library. The epitopes identified belonged predominantly to class I, followed by class IV and class III. Sera from mice immunized with either Spike or RBD of the ancestral strain exhibited similar epitope profiles, which were also similar to the profile of sera from mice immunized with Omicron RBD. In the Omicron RBD sera, we primarily detected epitopes at sites conserved between WT and Omicron RBD, dominated by class I epitope residues. Interestingly, the sera from BALB/c mice also contained antibodies targeting a cryptic epitope with conserved residues in all the current VOCs. The cryptic epitope, a class V epitope, was missed in the previous DMS epitope mapping scans until it was recently identified ^23,35–37^. The enrichment of the tested sera in antibodies targeting this epitope could help explain the cross-reactivity of these sera, as demonstrated by their neutralization potency against both B.1 and BA.1. The class V epitope targeting antibodies were also elicited in sera from individuals who received two doses of ChAdOX nCoV-19 vaccine and undergone subsequent breakthrough infections with SARS-CoV-2 variants ^24^. The K18-hACE2 mouse model expressing human (h)ACE2 is susceptible to SARS-CoV-2 infection and recapitulates the severity of the human disease. In this study, humanized mice were immunized with BA.1 RBD. The results showed that Omicron vaccination or infection elicited ACE2 competing antibodies and not class V-targeted antibodies. However, before the class V epitope was identified, there was a reference to it as the “E465 patch’’, and human sera from individuals infected with the ancestral SARS-CoV-2 strain showed the residues around the E465 patch escape from neutralization by the sera ^34^, which agrees with the results that we obtained from BALB/c mice. Overall, we do not have a comprehensive explanation for the difference observed between the epitopes targeted in K18-hACE2 and BALB/c mice. However, it is worth noting that the germline repertoire of the two strains is quite different. To infer the immunoglobulin genes of BALB/c mice, long-read SMRT sequencing was primarily used to amplify VDJ-C sequences from F1 (BALB/c x C57BL/6) hybrid animals. The study led to the identification of 278 germline IGHV alleles. Of these, 169 alleles were absent in the C57BL/6 genome reference sequence ^38,39^.

Sera from K18-hACE2 mice immunized with RBD from the ancestral strain exhibited a similar epitope profile to that of the BALB/c mice, enriched with class I, II, and IV and with less enrichment of class III. In contrast, sera from K18-hACE2 mice immunized with RBD from the Omicron BA.1 strain showed a different epitope profile relative to sera from K18-hACE2 mice immunized with RBD from the ancestral strain and sera from BALB/c mice immunized with the Omicron BA.1 strain. Sera of K18-hACE2 mice immunized with Omicron RBD focus on class I and class II epitopes. Lower targeting of the class V cryptic epitope was observed in K18 mice sera compared to BALB/c sera.

RBD is the major target for the neutralizing response. Using epitope mapping to analyse the regions of RBD targeted by different vaccine formulations or infections by SARS-CoV-2 WT and other VOCs can improve vaccine design and aid in developing an improved vaccine that elicits a cross-reactive immune response targeting conserved regions in the RBD.

Rapid, efficient, accurate, and inexpensive epitope mapping methods are essential for vaccine development and understanding humoral responses and viral evolution during pandemics or epidemics. The studies described here demonstrate that charged scanning mutagenesis libraries coupled with yeast surface display and deep sequencing can fulfil the requirements for high-resolution mapping tools that are easy to construct and analyse, and are time and cost-effective and efficient for rapid and high throughput screening. Another significant advantage of low diversity libraries is that they can be utilized for mammalian display, reverse genetics, and other approaches where the construction, characterization, and sequencing of complete saturation mutagenesis libraries can be challenging.

## Materials and Methods

### Library design and construction

SARS-CoV-2 RBD (332-532) was selected as a candidate for epitope mapping. A total of 124 residues were selected based on their solvent accessibilities extracted from the crystal structure of RBD with the ACE2 receptor (6M0J). All residues with accessibility ≥15% were selected for mutagenesis. Initially, the cutoff was set to 5%, but some charged mutations with a less than 15% cutoff showed an inconsistent effect on the protein’s expression. Hence, the cutoff was increased to 15%. Only one mutation with a cutoff of >15% showed reduced expression, Y453D, which was excluded from the library. All residues with accessibility ≥15% were mutated to either Aspartate or, if the residue was Aspartate, Asparagine or Glutamate, the residue was mutated to the positively charged residue, Arginine. The RBD sequence was cloned into the pUC57 plasmid with flanking regions of around 100 bp from the pETcon plasmid (Addgene plasmid # 41522). RBD-pUC57 plasmid was used as a template to individually introduce charged mutations at the selected positions using overlap PCR. Three unique primers were designed for each position, two mutagenesis primers and one pUC57 vector-specific reverse primer containing a 6-nucleotide (6N) long barcode with a known sequence and a 30 bp flanking sequence from the pETcon plasmid. A common pUC57 vector-specific forward primer with a 30 bp flanking sequence from the pETcon plasmid was designed. Briefly, the forward mutagenesis primer and the vector-specific barcoded reverse primer were used to generate the first PCR fragment. Similarly, the reverse mutagenesis primer and the vector-specific common forward primer were used to generate the second PCR fragment. The template used in the PCR to generate the two fragments was RBD-pUC57 plasmid at a concentration of 5 ng per reaction. The PCR reaction volume was 20 µL, and the used concentration of the primers was 0.25 µM. The PCR conditions used were: initial denaturation at 95 ℃ for 3 min, 7 cycles of (denaturation at 95 ℃ for 30 s, annealing at 60 ℃ for 30 s, and extension at 72 ℃ for 1 min), final extension at 72 ℃ for 5 min. By the end of the first round of PCR, both generated fragments have an overlapping region of around 24 bases. A second round of PCR was used to join the two fragments using a new pair of common, vector-specific, forward and reverse primers that bind the 30 bp pETcon flanking sequence in the pUC57 vector-specific forward primer. The template used in the second round of PCR was 1 µL (∼ 5 ng) of each of the two fragments generated by the first round of PCR without further purification. The reaction volume was 20 µL, and the primer concentration was 1 µM. The PCR conditions used were: initial denaturation at 95 ℃ for 3 min, followed by 15 cycles of denaturation at 95 ℃ for 30 s, annealing at 60 ℃ for 30 s, extension at 72 ℃ for 1 min, and a final extension at 72 ℃ for 5 min. After joining the two PCR fragments individually for each position, equal amounts of PCR joined product from each position were pooled and gel-purified.

The purified PCR pooled product was transformed into the yeast strain *EBY100* of *Saccharomyces cerevisiae* via electroporation using two fragment recombination along with NdeI and BglII (New England Biolabs) double digested pETcon plasmid, as described ^40^. Briefly, a single colony of EBY100 was inoculated and grown overnight in YPD media (10 g/L yeast nitrogen base, 20 g/L Peptone and 20 g/L Glucose) on a platform shaker at 225 rpm and 30 °C. The following day, a secondary culture of 100 mL YPD was inoculated from the overnight culture to a final OD_600_ of 0.3 and incubated at 30 °C and 225 rpm for about 5 hours until OD_600_ reached approximately 1.6.

The yeast cells were then harvested by centrifugation at 3000 rpm for 3 min, the media was removed, and the yeast cell pellet was washed twice with 50 mL ice-cold water and once with 50 mL of ice-cold electroporation buffer (1 M Sorbitol / 1 mM CaCl2). The yeast cell pellet was resuspended in 20 mL of 0.1 M LiAc and 10 mM DTT and incubated for 30 min at 30 °C and 225 rpm. The yeast cells were collected by centrifugation at 3000 rpm for 3 min, followed by one wash with 50 mL ice-cold electroporation buffer and then resuspension in around electroporation buffer to reach a final volume of 800 µL, which is sufficient for two transformation reactions. The purified PCR product (0.5 µg) along with 1 µg of double-digested pETcon vector were added to 400 µL yeast cells in the electroporation reaction, mixed by tapping, and transferred to a pre-chilled BioRad GenePulser cuvette (0.2 cm) (Bio-Rad, Hercules, CA, USA). The cells were electroporated at 2.5 kV for 3.5 ms and were transferred immediately to 8 mL of a 1:1 mix of 1 M sorbitol: YPD media and incubated for 1 hour at 30 °C and 225 rpm. The transformed yeast cells were collected by centrifugation and resuspended in 100 mL of SDCAA media (20 g/L dextrose, 6.7 g/L Difco yeast nitrogen base without amino acids, 5 g/L Bacto casamino acids, 14.7 g/L sodium citrate dihydrate, and 4.29 g/L citric acid monohydrate), incubated at 30 °C and 225 rpm overnight and then stored at −80 °C.

### Library sequencing

The transformed yeast surface-displayed aspartate barcoded library was deep sequenced to confirm the correct linkage between barcodes and mutations. A series of forward primers was designed to span the entire gene, with 100-base-pair separating each consecutive pair of primers. A common reverse primer binding downstream of the barcode region was designed. PCR reactions were carried out for each forward primer coupled with a common reverse primer in a 50 µL reaction and using 5 µL of the plasmid isolated from the yeast culture as a template. The PCR reactions were carried out for 20 cycles. The primers were used at a concentration of 0.5 µM and the PCR reaction was performed using the following conditions: initial denaturation at 95 ℃ for 3 min, 20 cycles of denaturation at 95 ℃ for 30 s, annealing at 60 ℃ for 30 s, and extension at 72 ℃ for 1 min, followed by final extension at 72 ℃ for 5 min. The PCR yielded fragments varying in length, ranging from the full gene down to 150 base pairs, with a 100-base-pair difference between each consecutive fragment. These fragments were gel purified, pooled in equimolar ratio and submitted for deep sequencing on an Illumina NovaSeq 150 PE. Fragments from the same variant shared identical barcodes, enabling their clustering and assembly to reconstruct the complete gene sequence.

### Mice immunization

The immunization studies were performed with mice species received after authorization from the Institutional Animal Ethics Committee (CAF/ETHICS/887/22). These experiments were carried out at the Central Animal Facility (CAF) of the Indian Institute of Science (IISC), adhering to CPCSEA and ARRIVE guidelines. In total, either female BALB/c mice aged 7-8 weeks (n=5) or K18-hACE-2 expressing C57BL/6 transgenic mice aged 8-9 weeks (n=6, comprising 3 males and 3 females) were used for immunization. A pre-bleed serum sample was collected one day prior to the immunization. The initial immunization took place on day zero, where 20 µg of each PBS-buffered protein (either stabilized RBD derived from Wuhan or omicron BA.1 SARS CoV-2 or soluble stabilized Spike ectodomain derived from Wuhan strain) was administered in conjunction with Sepivac SWE™ adjuvant (a squalene-in-water emulsion) in a 1:1 ratio, delivered in a total volume of 100 µL via intramuscular injection into both hindlimbs of the mice (50 µL per limb). Prime bleed sera were collected 14 days following the immunization. A booster immunization was administered 7 days later, utilizing the same dosage as the initial immunization, with boost sera collected on day 35.

### Pseudovirus generation and neutralization assay

Pseudoviruses were prepared in HEK-293T cells by transient transfection of Spike and HIVΔenv-nLuc backbone constructs as described previously ^41^. The neutralization assays were performed using 293T-hACE2-TMPRSS2 and frozen stocks of pseudoviruses. Briefly, sera were serially diluted in growth media, and pseudovirus was incubated for 1 hour, after which 1.5 × 10^4^ cells/well were added to make the final volume up to 200µL. The plate was incubated for 48 hours at 37 °C in a humidified incubator with 5% CO₂. The relative luminescence was measured by using the Nano-Glo luciferase assay system (Promega Inc., Catalog No. N1120) in a Cytation-5 multimode plate reader (BioTech Inc.). The infection control without serum was considered as 100% infection, and the percent reduction in infection (i.e., neutralization) was calculated accordingly. The serum inhibitory dilution (ID50) required for 50% neutralization of the pseudovirus was calculated using GraphPad Prism 8 software.

### Sera screening and flow cytometry

The yeast library was revived from −80 °C stock and grown overnight in 25 mL SDCAA media to a final OD_600_ of ∼ 0.3 at 30 °C, 250 rpm. The next day, the primary overnight culture was diluted in fresh 10 mL of SDCAA media to an OD_600_ of 0.5 and grown for 5 h at 30 °C before induction. The secondary culture was induced in 10 mL of SGCAA media (20 g/L galactose, 6.7 g/L Difco yeast nitrogen base without amino acids, 5 g/L Bacto casamino acids, 14.7 g/L sodium citrate dihydrate, and 4.29 g/L citric acid monohydrate) and incubated at 20 °C for 16 h. For library expression and binding experiments, cells from 1 OD of yeast culture were pelleted by centrifugation at 10,000 rpm for 1 min and washed twice with 250 µL of labelling buffer consisting of 1x phosphate buffered saline (PBS) and 0.5% bovine serum albumin (BSA) and incubated in 200 µL reaction with diluted chicken anti-c-Myc antibody (1:300) (Immunology Consultants Laboratory, Inc., Oregon, USA, CMYC-45A) for expression and ACE2/mAbs/sera for binding for 1 hr at 4 °C and 350 rpm. The labelled cells were washed thrice in labelling buffer and then incubated for 20 min at room temperature without shaking in 200 µL reactions with the secondary antibodies, anti-chicken-Alexa Fluor-488 (Thermo Fischer Scientific, A-11039) (1:300 dilution) for expression and anti-human/mouse-Alexa Fluor-633 (Thermo Fischer Scientific, A-21091 and A-21063) (1:1200 dilution) for binding. The labelled cells were resuspended in 1 mL labelling buffer for FACS sorting.

The immune sera were heat-inactivated at 56 °C for 1 h, centrifuged, and the supernatant was transferred to a fresh tube before use. Each serum sample was titrated on WT-RBD displayed on the yeast surface to obtain a titration curve. The sera were serially diluted and incubated with 0.1 OD_600_ of cells displaying WT-RBD for 1h, and the binding was detected by anti-human-Alexa Fluor 633 on a flow cytometer. From the titration curve, a specific serum dilution was selected and subjected to epitope mapping with the charged scanning library.

Using a BD FACS Aria III, the yeast library was sorted into different gates from the 1D histograms based on Alexa Fluor-488 for expression or Alexa Fluor-633 for binding. The first gate contains the unlabeled cells and the rest of the gates were distributed to span the whole binding/expression MFI range. From each bin, 100,000 cells were collected in 1 mL of SDCAA media and grown at 30 ℃ and 250 rpm in 10 mL of SDCAA media until saturation.

### Illumina sequencing

The sorted samples were grown for 24 hours in 10 mL of SDCAA media at 30 °C, 250 RPM, and the plasmids were isolated from yeast cells using Longlife™ Zymolyase® (G Biosciences). Briefly, the yeast cells were collected by centrifugation at 3000 rpm for 5 min and then resuspended in a 150 µL plasmid resuspension buffer containing RNase. Zymolyase (10 units) was added to the resuspended cells, and samples were incubated for 5 hours at 37 ℃ and 180 rpm. After 5 hours at 37 ℃, the resuspended cells were frozen at −20 ℃ for 6 hours, and then an additional 150 µL of resuspension buffer was added, followed by 200 µL of Lysis buffer. After incubation with lysis buffer for 4 min, 490 µL of neutralization buffer was added and the cell lysate was centrifuged at 15,000 rpm for 15 min. An equal volume of Isopropanol was added to the clear supernatant, and this was passed through a silica column (GeneJET, ThermoScientific). The column was then washed with 750 µL of wash buffer. The column was subjected to dry spin for 2 min at 15,000 rpm before the DNA was eluted in 50 µL autoclaved Milli Q water. The isolated plasmids were used as a template for PCR along with forward and reverse primers flanking the barcode region, to yield a PCR product with a size of 150 bp. The PCR primers were designed to amplify the barcode region only (150 bp) and in a way such that for both the forward and reverse end reads, the first 3 bases are *NNN* followed by **6 bases of unique sequence tag** (**MID**) and the 18 to 21 bases of the primer sequence <u>complementary to the gene</u>, for example (*NNN***TGTGGG**<u>ACTGCGGGGGCCTCGAGG</u>). Each forward MID sequence represents a particular serum, while each reverse MID sequence represents a specific gate. The PCR for all the gates from three sorted sera samples was carried out using Phusion polymerase for 15 cycles. Following agarose gel electrophoresis and gel band purification, the PCR products were quantified by NanoDrop 2000c (Thermo Fisher Scientific, Waltham, MA, USA) and pooled in equimolar ratio (∼100 ng of PCR product from each sample), before submitting for sequencing on the 150 PE NovaSeq platform.

### Processing of deep sequencing data for library validation

The initial step involved assembling paired-end reads using the PEAR v0.9.6 (Paired-End Read Merger) tool ^42^. Subsequently, the reads were segregated based on distinct Molecular Identifiers (MIDs) for each gate. The integrity of both forward and reverse primers was individually verified before merging the files containing both types of reads.

A “quality filtering” process ensued, removing terminal “NNN” residues and excluding reads lacking the relevant MID and/or primers. Reads with mismatched MIDs were also discarded. The reads were then categorized based on the gates associated with expression and binding (comprising 16 gates) for all forward primers. After removing “NNN” residues, the .fastq files were converted to .fasta files. Subsequently, the reads underwent further filtration, retaining only those with a minimum length of 118 bases.

The length-filtered reads were subjected to alignment with the wild-type (WT) sequence using the Water v6.4.0.0 program. ^43^ and reformatted accordingly. Barcodes were extracted from the reformatted alignment along with their corresponding read coordinates. These extracted barcodes were cross-referenced with a predefined list of 124 barcodes used in the experiments. The frequency of matched barcodes was calculated, and a table was generated, listing the matched barcodes and their corresponding reads within the gates across duplicate samples.

### Analysis of the deep sequencing reads

The deep sequencing of the barcodes from each bin was subjected to analysis as described previously. ^19^. We first normalized the reads of each mutant across different bins (Equation 3.1), then calculated the fraction (*Xi*) of each mutant *i* distributed across all the different bins (Equation 3.2). The reconstructed MFI for each mutant was calculated by summing the product of the mutant fraction (*Xi*) in a particular bin and the MFI of that bin (*Fi*) obtained from the FACS experiment across all the bins populated by that mutant (Equation 3.3). Finally, we normalized the expression and binding of each mutant with respect to the WT (Equation 3.4).

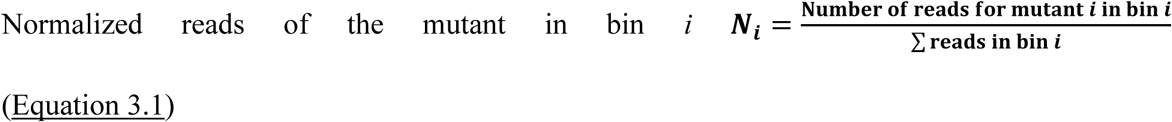

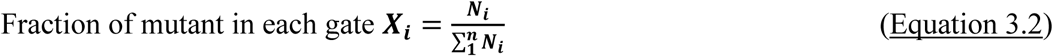

N*i*: Number of reads of mutant in bin *i*

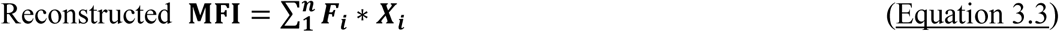

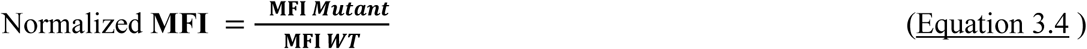

The mutants with Ratio, MFI_Ratio_ (Normalized MFI of expression: Normalized MFI of binding), one standard deviation higher than the mean MFI, which corresponds to a z-score of +1, were considered as interacting or epitope residues. The MFI_Ratio_ of ACE2 binding for all the library residues was plotted as a heatmap showing the identified interacting residues with MFI_Ratio_ higher than the rest of the residues in the library in red colour; the darker the red colour, the higher the escape from ACE2 binding exhibited by this mutant. The identified interacting residues, except for one, were indeed interacting residues as reported in the literature and confirmed by examining the RBD-ACE2 bound crystal structure. The charged scanning epitope mapping methodology precision and recall were calculated using the following formulas:

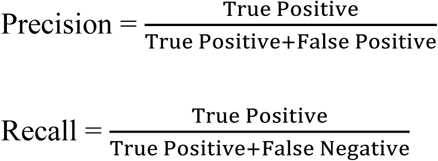

### ΔASA, Δ Depth, and interacting residues Calculations

Crystal structures of RBD alone and ACE-2-bound RBD were used to identify RBD residues interacting with ACE2 (PDB ID: 6VXX and PDB ID: 6M0J). ΔASA for unbound and ACE2-bound RBD were calculated using NACCESS ^22^, as the difference between the solvent-accessible surface areas of RBD residues in its free form (PDB ID: 6VXX) and ACE2-bound form (PDB ID: 6M0J). Residues with ΔASA>0 were considered interacting residues. ΔDepth was calculated as the difference in residue depth between the RBD-ACE2 bond structure (6M0J) and the RBD unbound structure (8SGU).

### Statistical analysis

Unless mentioned otherwise, all experiments have been performed in duplicates (n=2).

## Author Contributions

R.V. and K.K. Conceptualization. K.K. performed the library constructions, epitope mapping experiments and the analyses. M.B. performed the DMS datasets analysis and processed the deep sequencing reads. C.D. and N.B. helped in methodology validation. R.S. performed the mouse immunization and ELISA. S.K. and R.P.R. performed the pseudovirus neutralization assay. A.A. helped in FACS sorting. R.V. and K.K. wrote the original draft with input from all the authors.

## Data availability

The data relevant to the figures in the paper have been made available within the article and in the supplementary information section. The deep sequencing data discussed in the present study have been deposited in the SRA database (accession number: PRJNA1207487).

## Supporting Information

The supplementary file contains Figure S1, which shows Cut-off validation for the MFI_Ratio_ of putative epitope residues; Figure S2 is a Structural representation of neutralizing antibody classes. Figure S3 shows the NGS validation. Table S1 shows the RBD residues selected for charge scanning mutagenesis. Table S2 shows the read distribution across the sequenced fragments with their quality filtered (>Q20) raw reads. Table S3 shows the characteristics of the interacting residues of SARS-CoV-2 RBD with the ACE2 receptor. Table S4 shows MFI_Ratio_ for the epitopes identified in this study for the various polyclonal sera.

