## Supporting file for "Charged Scanning Mutagenesis as a High Throughput Approach for Epitope Mapping"

**Table S1.** SARS-CoV-2 RBD residues selected for charge scanning mutagenesis. The residues are solvent-exposed with accessibility ≥15%.

| Mutated Positions of RBD | | | | | | | | |
|---|---|---|---|---|---|---|---|---|
| I332 | N354 | S373 | R403 | N439 | S459 | A475 | Q493 | H519 |
| T333 | R355 | S375 | D405 | N440 | N460 | G476 | S494 | A520 |
| N334 | K356 | T376 | R408 | L441 | K462 | S477 | G496 | P521 |
| L335 | R357 | K378 | G413 | K444 | P463D | T478 | Q498 | A522 |
| P337 | S359 | Y380 | T415 | V445 | F464D | P479 | P499 | T523 |
| G339 | N360 | G381 | G416 | G446 | E465 | N481 | T500 | P527 |
| E340 | V362 | S383 | K417 | N448 | R466 | G482 | N501 | K528 |
| N343 | D364 | P384 | D420 | Y449 | D467 | V483 | G502 | K529 |
| A344 | S366 | T385 | Y421 | N450 | I468 | E484 | V503 | S530 |
| T345 | V367 | K386 | K424 | L452 | S469 | G485 | G504 | T531 |
| R346 | Y369 | L387 | D427 | L455 | T470 | F486 | Y505 | N532 |
| A348 | N370 | D389 | D428 | F456 | E471 | N487 | E516 | G533 |
| Y351 | S371 | L390 | T430 | R457 | Y473 | Y489 | L517 | |
| A352 | A372 | Y396 | N437 | K458 | Q474 | F490 | L518 | |

**Table S2.** Reads distribution across the sequenced fragments with their quality filtered (Phred score > Q20) raw reads.

| Mutant_Barcode | Frag 6 | Frag 5 | Frag 4 | Frag 3 | Frag 2 | Frag 1 | TOTAL |
|---|---|---|---|---|---|---|---|
| I332D_CTTGAT | 5 | | | | | | 5 |
| T333D_TTTTTT | 27 | | | | | | 27 |
| N334R_GGTTTT | 38 | | | | | | 38 |
| L335D_GTGTTT | 8 | | | | | | 8 |
| P337D_TGGTTT | 20 | | | | | | 20 |
| G339D_ACGTTT | 23 | | | | | | 23 |
| E340R_CAGTTT | 19 | | | | | | 19 |
| T345D_TCCTTT | 3 | | | | | | 3 |
| R346D_CCAATT | 2 | | | | | | 2 |
| A348D_AAAATT | 18 | | | | | | 18 |
| Y351D_GTTTGT | 20 | | | | | | 20 |
| A352D_TGTTGT | 8 | | | | | | 8 |
| N354R_GGTGGT | 21 | | | | | | 21 |
| R355D_CCTGGT | 9 | | | | | | 9 |
| K356D_AATGGT | 27 | | | | | | 27 |
| S359D_TGGGGT | 15 | | | | | | 15 |
| N360R_ACGGGT | 17 | | | | | | 17 |
| V362D_CAGGGT | 7 | | | | | | 7 |
| D364R_TAAGGT | 31 | | | | | | 31 |
| S366D_ATTCGT | 1 | | | | | | 1 |
| V367D_CGTCGT | 7 | | | | | | 7 |
| Y369D_GCTCGT | 17 | 22 | | | | | 39 |

| | | | | | | | |
|---|---|---|---|---|---|---|---|
| N370R_TATCGT | 24 | 40 | | | | | 64 |
| S371D_TCGCGT | 11 | 22 | | | | | 33 |
| A372D_GAGCGT | 25 | 26 | | | | | 51 |
| S373D_GTCCGT | 13 | 16 | | | | | 29 |
| S375D_TGCCGT | | 40 | | | | | 40 |
| T376D_ACCCGT | | 33 | | | | | 33 |
| K378D_CACCGT | | 11 | | | | | 11 |
| Y380D_CAAAGT | | 28 | | | | | 28 |
| G381D_AGTTCT | | 14 | | | | | 14 |
| S383D_ATGTCT | | 49 | | | | | 49 |
| P384D_TTCTCT | | 34 | | | | | 34 |
| T385D_GTATCT | | 24 | | | | | 24 |
| K386D_CAATCT | | 31 | | | | | 31 |
| L387D_CTGGCT | | 33 | | | | | 33 |
| D389R_AGGGCT | | 39 | | | | | 39 |
| L390D_TTAGCT | | 50 | | | | | 50 |
| Y396D_GGAGCT | | 25 | | | | | 25 |
| R403D_AAAGCT | | 49 | 112 | | | | 161 |
| D405R_CCTCCT | | 37 | 83 | | | | 120 |
| R408D_TCCCCT | | 23 | 59 | | | | 82 |
| G413D_CGACCT | | | 65 | | | | 65 |
| T415D_GGGACT | | | 173 | | | | 173 |
| G416D_ATCACT | | | 126 | | | | 126 |
| K417D_GAAACT | | | 34 | | | | 34 |
| D420R_AGGTAT | | | 14 | | | | 14 |
| Y421D_TGCTAT | | | 3 | | | | 3 |

| | | | | | | | |
|---|---|---|---|---|---|---|---|
| K424D_AAATAT | | | 24 | | | | 24 |
| D427R_AGTGAT | | | 7 | | | | 7 |
| D428R_CGGGAT | | | 86 | | | | 86 |
| T430D_CCCGAT | | | 73 | | | | 73 |
| N437R_GTTCAT | | | 8 | 7 | | | 15 |
| N439R_GGGCAT | | | 78 | 61 | | | 139 |
| N440R_CCGCAT | | | 83 | 50 | | | 133 |
| L441D_AATAAT | | | 145 | 110 | | | 255 |
| K444D_CTCAAT | | | 100 | 86 | | | 186 |
| V445D_GACAAT | | | | 69 | | | 69 |
| G446D_GTTTTG | | | | 319 | | | 319 |
| N448R_GGGTTG | | | | 76 | | | 76 |
| Y449D_ATCTTG | | | | 82 | | | 82 |
| N450R_TAGATG | | | | 112 | | | 112 |
| L452D_ACAATG | | | | 108 | | | 108 |
| L455D_CCTTGG | | | | 184 | | | 184 |
| F456D_TAATGG | | | | 260 | | | 260 |
| R457D_CATGGG | | | | 74 | | | 74 |
| K458D_GGGGGG | | | | 24 | | | 24 |
| S459D_CCCCGG | | | | 106 | | | 106 |
| N460R_AGGAGG | | | | 12 | | | 12 |
| K462D_GGAAGG | | | | 59 | | | 59 |
| P463D_AAAAGG | | | | 70 | | | 70 |
| F464D_GCCCCG | | | | 67 | | | 67 |
| E465R_AAAGAG | | | | 26 | | | 26 |
| R466D_GTTAAG | | | | 162 | | | 162 |

| | | | | | | | |
|---|---|---|---|---|---|---|---|
| D467R_CGCAAG | | | | 75 | | | 75 |
| I468D_AGAAAG | | | | 77 | | | 77 |
| S469D_GAAAAG | | | | 23 | 74 | | 97 |
| T470D_AGTTTC | | | | 17 | 39 | | 56 |
| E471R_CCCTTC | | | | 11 | 25 | | 36 |
| Y473D_TCGGTC | | | | 21 | 63 | | 84 |
| Q474D_ACCGTC | | | | 57 | 191 | | 248 |
| A475D_TTTCTC | | | | 16 | 70 | | 86 |
| G476D_TCCCTC | | | | 32 | 106 | | 138 |
| S477D_CGACTC | | | | 55 | 234 | | 289 |
| T478D_GTTATC | | | | | 206 | | 206 |
| P479D_TTGATC | | | | | 500 | | 500 |
| N481R_GCCATC | | | | | 247 | | 247 |
| G482D_GAAATC | | | | | 318 | | 318 |
| V483D_ATTTGC | | | | | 383 | | 383 |
| E484R_AAATGC | | | | | 427 | | 427 |
| G485D_CGGGGC | | | | | 92 | | 92 |
| F486D_CCCGGC | | | | | 190 | | 190 |
| N487R_ACAGGC | | | | | 167 | | 167 |
| Y489D_CCGCGC | | | | | 207 | | 207 |
| F490D_TCACGC | | | | | 277 | | 277 |
| Q493D_TGGAGC | | | | | 423 | | 423 |
| S494D_CCTTCC | | | | | 438 | | 438 |
| G496D_GACTCC | | | | | 250 | | 250 |
| Q498D_GTTGCC | | | | | 259 | | 259 |
| P499D_CCGGCC | | | | | 532 | | 532 |

| | | | | | | | |
|---|---|---|---|---|---|---|---|
| T500D_CTTCCC | | | | | 527 | | 527 |
| N501R_GCGCCC | | | | | 595 | | 595 |
| G502D_CAACCC | | | | | 676 | 1638 | 2314 |
| V503D_AGGACC | | | | | 576 | 2232 | 2808 |
| G504D_AAAACC | | | | | 360 | 1272 | 1632 |
| Y505D_GGGTAC | | | | | 415 | 1929 | 2344 |
| E516R_GCCTAC | | | | | | 534 | 534 |
| L517D_TTTGAC | | | | | | 1251 | 1251 |
| L518D_TCCGAC | | | | | | 2540 | 2540 |
| H519D_ACCCAC | | | | | | 2828 | 2828 |
| A520D_CAAAAC | | | | | | 2490 | 2490 |
| P521D_ACCTTA | | | | | | 2113 | 2113 |
| A522D_CCCGTA | | | | | | 2912 | 2912 |
| T523D_TTTATA | | | | | | 2120 | 2120 |
| S530D_TCCCGA | | | | | | 3731 | 3731 |
| T531D_GGGAGA | | | | | | 2734 | 2734 |
| N532R_AAACCA | | | | | | 3445 | 3445 |
| G533D_TTTTAA | | | | | | 4467 | 4467 |

**Table S3.** Interacting residues of SARS-CoV-2 RBD with the ACE2 receptor based on the ΔASA calculation. ΔASA is the difference between the solvent-accessible surface area of the RBD residues in the free form and the ACE2-bound form. Key interacting residues with ΔASA ($\geq 0$ Å$^2$) are presented in the table. Delta accessibility (ΔASA) values were calculated based on the crystal structure of ACE2-bound RBD (6M0J) and delta Depth (ΔDepth) values were calculated from subtracting the residue depth in the ACE2 bound RBD structure (6M0J) from the residue depth in the RBD unbound structure (8SGU) (Tan et al., 2011). $MFI_{Ratio}$ was calculated from the deep sequencing reads analysis as explained in the methods. (NA no reads). All residues with $MFI_{Ratio}$ > $MFI_{Ratio}$+1 were assumed to be epitope residues as described in Figure S1.

| Position | Residue | ΔASA -All atom (Å$^2$) | ΔDEPTH (Å) | $MFI_{Ratio}$ |
|---|---|---|---|---|
| 403 | ARG | 0.8 | 0.4 | 2.17 |
| 417 | LYS | 17.6 | 0.21 | 1.51 |
| 445 | VAL | 6.2 | 0.29 | 1.45 |
| 446 | GLY | 22.7 | 0.17 | 1.37 |
| 449 | TYR | 19.8 | −0.01 | 1.62 |
| 453 | TYR | 13.2 | −1.78 | NA |
| 455 | LEU | 29.2 | −3.32 | 3.25 |
| 456 | PHE | 27.6 | −2.14 | 2.19 |
| 473 | TYR | 3.5 | −0.31 | 3.83 |
| 475 | ALA | 23.3 | −2.22 | 3.65 |
| 476 | GLY | 31.7 | −0.23 | 1.32 |
| 484 | GLU | 9.8 | 0.01 | 1.38 |
| 486 | PHE | 60 | −0.36 | 2.13 |
| 487 | TYR | 34.1 | −1.46 | 2.58 |
| 489 | ASN | 40.2 | −2.79 | 2.71 |
| 493 | GLY | 38 | −1.25 | 2 |
| 496 | GLN | 21.5 | −3.84 | 3.28 |
| 498 | GLN | 39.6 | −2.28 | 1.94 |
| 500 | THR | 66.9 | −0.83 | 2.18 |
| 501 | GLY | 28.1 | −4.82 | 3.2 |
| 502 | ASN | 71.4 | −1.3 | 2.42 |
| 503 | VAL | 7 | 0.03 | 1.27 |
| 505 | TYR | 49.3 | −1.54 | 2.53 |

**Table S4.** $MFI_{Ratio}$ for the epitopes identified in this study for the various polyclonal sera. ( _ denotes non-epitope residue).

| Residue # | MFI Ratio | | | | |
|---|---|---|---|---|---|
| | BALB/c mice | | | K18-Tg mice | |
| | B.1 Spike | B.1-RBD | BA.1-RBD | B.1-RBD | BA.1-RBD |
| 346 | __ | __ | __ | __ | 1.16 |
| 351 | 1.58 | 1.44 | 1.55 | 1.44 | __ |
| 355 | 1.39 | 1.22 | 1.39 | __ | __ |
| 356 | 1.25 | 1.23 | 1.31 | 1.19 | __ |
| 357 | __ | __ | __ | __ | 1.07 |
| 359 | __ | __ | __ | __ | 1.06 |
| 364 | 1.34 | 1.25 | 1.27 | | __ |
| 376 | __ | __ | __ | 1.19 | __ |
| 378 | __ | __ | __ | | 1.09 |
| 380 | __ | __ | 1.32 | __ | __ |
| 381 | 1.34 | 1.30 | 1.34 | 1.27 | __ |
| 387 | 1.27 | 1.27 | __ | 1.22 | __ |
| 390 | __ | 1.23 | __ | __ | __ |
| 396 | __ | 1.25 | __ | __ | __ |
| 403 | 1.30 | 1.31 | 1.32 | 1.24 | __ |
| 413 | __ | 1.22 | __ | __ | __ |
| 416 | 1.66 | 1.44 | 1.54 | 1.37 | __ |
| 421 | 1.62 | 1.56 | 1.37 | 1.56 | __ |
| 424 | 1.32 | __ | 1.44 | 1.22 | __ |
| 430 | 1.50 | 1.41 | 1.41 | 1.35 | __ |
| 437 | 1.45 | 1.34 | 1.23 | 1.21 | __ |
| 448 | __ | __ | __ | __ | 1.24 |
| 450 | __ | __ | __ | __ | 1.11 |
| 452 | __ | __ | __ | __ | 1.09 |
| 456 | __ | __ | __ | __ | 1.40 |
| 457 | 1.37 | 1.27 | 1.30 | 1.27 | 1.12 |
| 459 | __ | __ | __ | __ | 1.16 |
| 462 | __ | __ | __ | __ | 1.12 |
| 463 | 1.23 | 1.20 | __ | __ | __ |
| 464 | 1.25 | 1.28 | __ | __ | __ |

| | | | | | |
|---|---|---|---|---|---|
| 465 | 1.37 | 1.26 | 1.32 | 1.18 | __ |
| 466 | 1.55 | 1.45 | 1.52 | 1.30 | __ |
| 467 | 1.50 | 1.31 | 1.51 | 1.28 | __ |
| 468 | __ | __ | 1.45 | __ | 1.14 |
| 469 | __ | __ | __ | __ | 1.12 |
| 470 | __ | __ | __ | 1.20 | 1.08 |
| 473 | __ | __ | __ | 1.22 | __ |
| 475 | __ | __ | __ | 1.19 | __ |
| 518 | __ | __ | __ | __ | 1.06 |
| 522 | __ | __ | __ | __ | 1.07 |

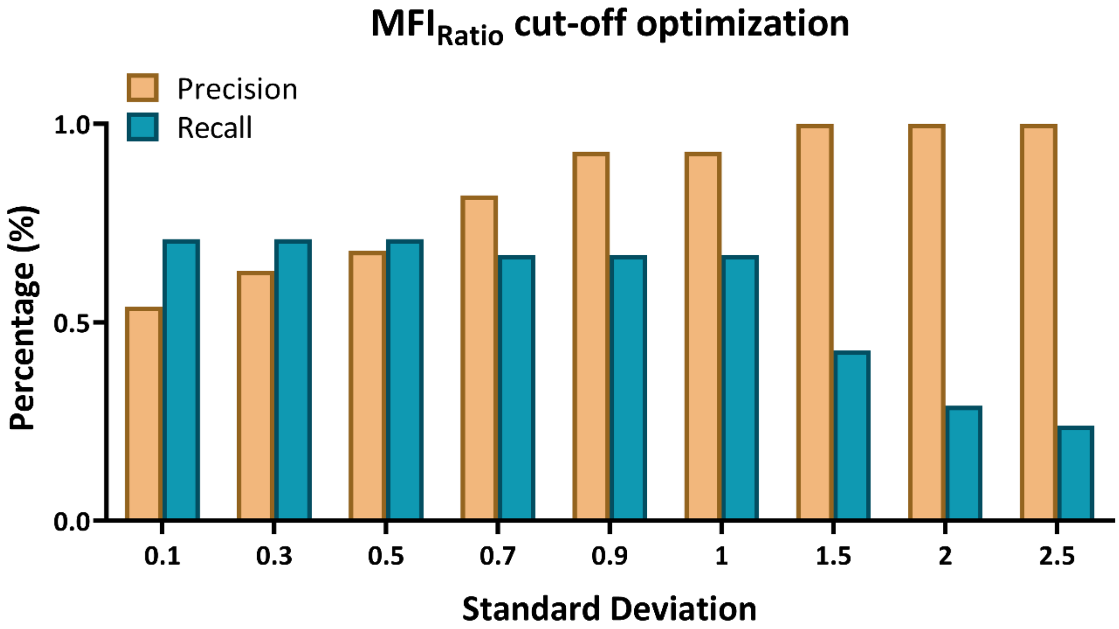


**Figure S1. Optimization of cut-off values for identification of putative RBD residues involved in ACE2 binding.** X is a variable, and SD is the standard deviation of the distribution of $MFI_{Ratio}$ values. X was varied from 0.1 to 2.5. In each case, residues with MFIRatio greater than MFI+XSD were assumed to be a part of the RBD: ACE2 interface. The best precision and recall values, 93% and 65%, respectively, were obtained by using the cutoff of X=1. Thus, mutants with $MFI_{Ratio}$ one standard deviation higher than the mean MFI were considered as ACE2 interacting or epitope residues.

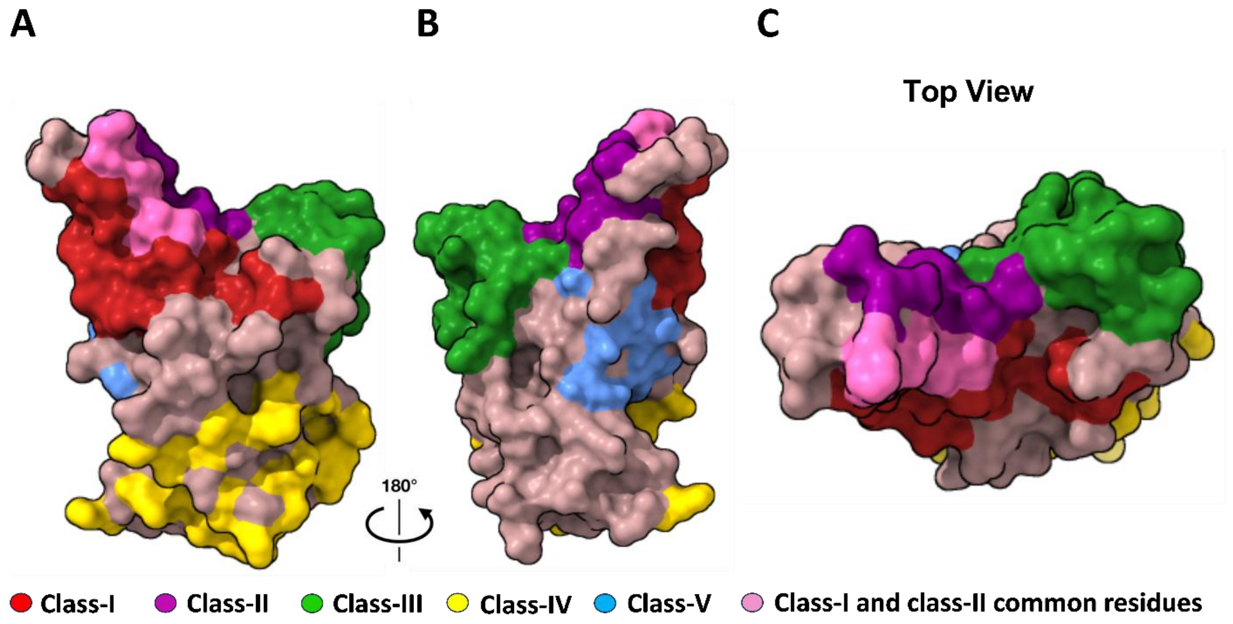


**Figure S2. Structural representation of neutralizing antibodies classes.** Five classes of neutralizing antibody epitopes are mapped on the structure of RBD (PDB: 6M0J). The class V epitope represents a cryptic epitope on the opposite face of the RBM from class I. (A) RBD inner face. (B) RBD outer face. (C) top view.

**A**

| Residues | $MFI_{Ratio}$ | Interacting |
|---|---|---|
| D428R | 1.36 | No |
| P463D | 1.57 | No |
| I468D | 1.26 | No |
| Y473D | 3.82 | Yes |
| A475D | 3.65 | Yes |
| G496D | 3.28 | Yes |
| N501R | 3.19 | Yes |
| Y505D | 2.52 | Yes |
| A520D | 1.30 | No |

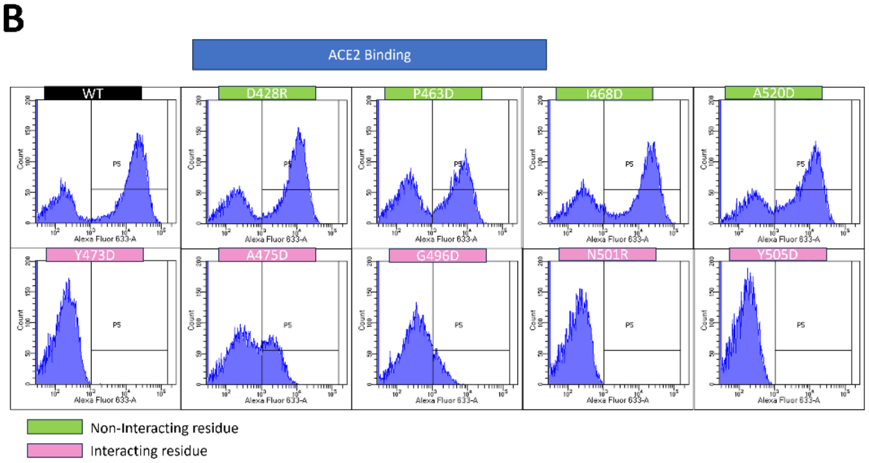


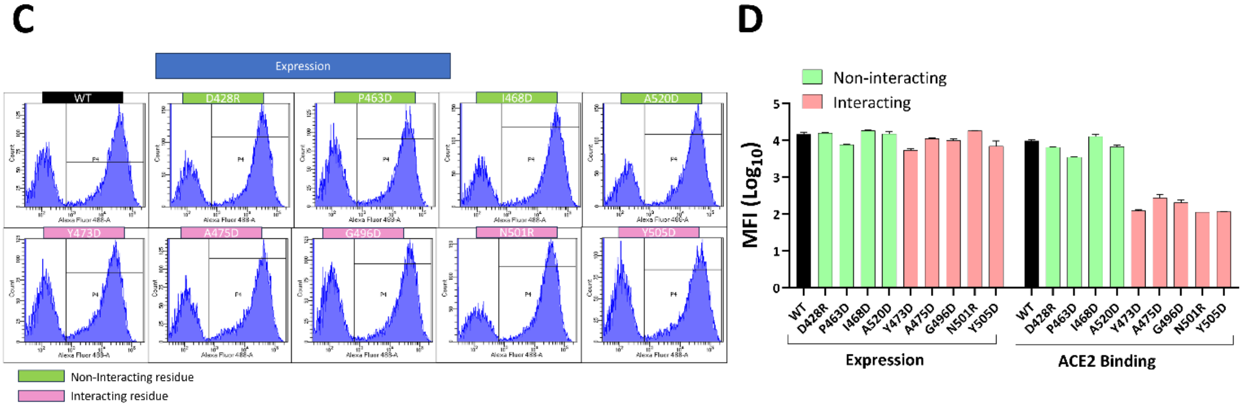


**Figure S3. Validation of the ACE2 interacting residues identified by NGS.** (A) Five and four residues identified in our NGS analysis as ACE2-interacting and non-interacting residues, respectively, in addition to the unmutated WT, were selected for validation by YSD. (B) Expression profiles on the yeast surface of the selected residues. (C) ACE2 binding profiles of the selected residues. (D) The bar graph represents the mean fluorescence intensity (MFI) of the expression and ACE2-binding of the residues selected for validation on the yeast surface.
